# Image-based Flow Simulation of Platelet Aggregates under Different Shear Rates

**DOI:** 10.1101/2023.02.22.529480

**Authors:** Yue Hao, Gábor Závodszky, Claudia Tersteeg, Mojtaba Barzegari, Alfons G. Hoekstra

## Abstract

Hemodynamics is crucial for the activation and aggregation of platelets in response to flow-induced shear. In this paper, a novel image-based computational model simulating blood flow through and around platelet aggregates is presented. The microstructure of aggregates was captured by two different modalities of microscopy images of *in vitro* whole blood perfusion experiments in microfluidic chambers coated with collagen. One set of images captured the geometry of the aggregate outline, while the other employed platelet labelling to infer the internal density. The platelet aggregates were modelled as a porous medium, the permeability of which was calculated with the Kozeny-Carman equation. The computational model was subsequently applied to study hemodynamics inside and around the platelet aggregates. The blood flow velocity, shear stress and kinetic force exerted on the aggregates were investigated and compared under 800 *s*^−1^, 1600 *s*^−1^ and 4000 *s*^−1^ wall shear rates. The advection-diffusion balance of agonist transport inside the platelet aggregates was also evaluated by local Péclet number. The findings show that the transport of agonists is not only affected by the shear rate but also significantly influenced by the microstructure of the aggregates. Moreover, large kinetic forces were found at the transition zone from shell to core of the aggregates, which could contribute to identifying the boundary between the shell and the core. The shear rate and the rate of elongation flow were investigated as well. The results imply that the emerging shapes of aggregates are highly correlated to the shear rate and the rate of elongation. The framework provides a way to incorporate the internal microstructure of the aggregates into the computational model and yields a better understanding of the hemodynamics and physiology of platelet aggregates, hence laying the foundation for predicting aggregation and deformation under different flow conditions.

**Author summary:** The initial step in the formation of an arterial thrombus is the rapid aggregation of the tiny blood particles called platelets. This process significantly influences the formation and structure of the resulting thrombi. The mechanical properties of the aggregates depend on their microstructure, which in turn is dictated by their interaction with the flow during formation. However, due to currently existing technological limitations, it is not possible to measure these interactions in sufficient detail experimentally. In this paper, an image-based computational model is proposed based on two different modalities of experimental images, that can complement the experiments and give detailed information on hemodynamics during the aggregation. The image sets are captured from whole blood perfused microfluidic chambers coated with collagen. One modality of images captured the shape of the aggregate outline with high contrast, while the other employed platelet labeling to infer the internal density. The platelet aggregates are considered as porous media in the simulations, informed by the images. This framework incorporates the internal microstructure of the aggregates into the computational model and yields a better understanding of the hemodynamics and physiology of platelet aggregates, hence laying the foundation for predicting aggregation and deformation under different flow conditions.

## Introduction

Platelet aggregation is the initial step of thrombus formation, that begins with platelet adhesion to the sites of vascular injury [1–3]. It is a multistep dynamic process involving distinct receptors and adhesive ligands such as GPIb*α* and *α*_IIb_*β*_3_ receptors, von Willebrand factor (VWF), fibrinogen and fibronectin [4]. Those receptor-ligand interactions regulate tethering, platelet-platelet cohesion and stabilization of the formed aggregates. Blood flow also plays a critical role in the process as distinct shear ranges lead to unique ways to form aggregates [4, 5]. Under low shear conditions, fibrinogen and integrin *α*_IIb_*β*_3_ are known to be the predominant factors of platelet aggregation while VWF and fibronectin have progressively increasing contribution to the aggregation with the increase of the flow shear rate [6].

To better understand the impact of the hemodynamics on the formation and mechanics of blood clots, several computational models were developed [7–12]. Since the porosity of the forming aggregate influences its formation [13–16], a growing number of computational studies considered the thrombus as a porous medium. Tomaiuolo et al. [17] established a two-dimensional computational model, in which the thrombus consists of two homogeneous porous parts. One with highly activated, densely packed platelets was named core and the other with lower levels of activation was named shell. Different permeabilities were assigned for the core and shell, respectively. They have shown that the solute transport behaviour changes with changes of the platelet packing density. Xu et al. [18] developed a similar novel two-dimensional multi-phase computational model to characterise the interplay between the main components of the clot. They showed that the shear force acting on the clot surface is affected by the internal structure of clots. Also, their results indicated that the porosity of the clots decreased as the flow shear rate increased.

The growing interest in the porous structure of thrombus or platelet aggregate has prompted measurements of their permeability [19, 20]. Experimental studies have demonstrated several ways to measure the permeability of clots [20–24]. However, owing to the significant differences in the setup, the composition of the flow and the heterogeneous nature of the formation, the experimental results of permeability usually correspond to unique conditions and are difficult to reuse in different scenarios. At present, the Kozeny-Carman formula [25, 26] is one of the most widely applied methods to estimate permeability. It approximates the permeability based on the volume fraction and geometric properties of the solid components that form the porous substance [14, 27].

Instead of considering the clot geometry as a simplified object such as half ellipse in the model, the geometry of the clot can be obtained from *in vivo* or *in vitro* imaging data, contributing to further studies on the microstructure of platelet aggregates [28]. For example, in [29], a two-dimensional model was developed to calculate the shear stress of the near-thrombus region using the fluorescent labeling images captured in mice. They quantitatively measured the thrombus dynamics in the early stages of hemostasis based on the images, but did not consider the internal structure of the thrombus. Taylor et al. [30] and Pinar et al. [31] examined the dynamics of the blood flow around the thrombus based on the *in vitro* experiments, but both of them considered thrombi as impermeable solids. In [32], the porous geometry of the clot was reconstructed from the experimental images. This setting is closer to reality compared to the impermeable solid and provides the prediction of the transport of inert solutes in the aggregates.

In this paper, an image-based method to capture the internal microstructure is proposed. Based on this method, a three-dimensional computational model is developed, that can simulate the blood flows through the formed platelet aggregates in a microchannel accurately. In this model, the platelet aggregates are considered as porous media and the microstructure of platelet aggregates is based on *in vitro* experiments of whole blood perfusion in microfluidic chambers coated with collagen. Two sets of images were used to capture the geometries and internal structures of the aggregates. The images without platelet labeling captured the geometry of the aggregate outline, while the images with platelet labeling are employed to infer the internal density of the aggregates and estimate the permeability. The computational model was subsequently applied to study the hemodynamics inside and around the platelet aggregates. The blood flow velocity, shear stress, kinetic forces of blood flow exerted on aggregates, flow elongational rate and advection-diffusion balance of agonist transport inside the platelet aggregates were investigated and compared under 800 *s*^−1^, 1600 *s*^−1^ and 4000 *s*^−1^ wall shear rates (WSRs). The proposed model incorporating the internal microstructure of the aggregates provides an accurate estimate of the hemodynamics of platelet aggregates, and thus lays the foundation for predicting further aggregation and deformation under different flow conditions.

## Materials and methods

### *In vitro* experiments

All the experiments were performed at the Laboratory for Thrombosis Research at KU Leuven. A schematic diagram of the *in vitro* experiments of platelet aggregates is shown in Fig 1. Fresh human blood was drawn into a citrated tube (3.8% sodium citrate, citrate:blood = 1:9) and mixed with MitoTracker Deep Red (Invitrogen, USA). Citrate chelates extracellular calcium, and without calcium coagulation cannot occur, hence fibrin cannot be formed. A temperature control (Thermo Fisher Scientific, US) was − used to ensure the blood was at a human-body temperature. The chip *µ* Slide VI 0.1 (Ibidi, Germany) was positioned under the microscope and connected to the test tube with blood and the pump via tubing. Before the experiments, the complete surface of the perfusion chamber was coated using 100 *µg/ml* Horm-collagen (Takeda, Austria) and stood overnight, resulting in a uniform distribution of collagen fibrils over the complete perfused area. The pump (Harvard Apparatus, US), which was on the other side of the chip, pulled the blood through the chamber with a preset WSR. After five minutes of blood perfusion, for every one micrometer in the z-direction, the images of cross-sections of the channel were captured under an Axio Observer Z1 inverted fluorescence microscope (Zeiss, Germany) using differential interference contrast (DIC) and fluorescence microscopy at 100x magnification. The perfusion time was chosen to ensure after which a stable clot was formed *in vitro*. The three-dimensional structure of platelet aggregates was reconstructed by stacking all the images over the z-direction later. Two types of images modality were captured for the platelet aggregates. One set of images captured the shape of the aggregate outline with non-labelled platelets (DIC images), while the other employed platelet labelling to infer the internal density (fluorescent images).

**Fig 1.**
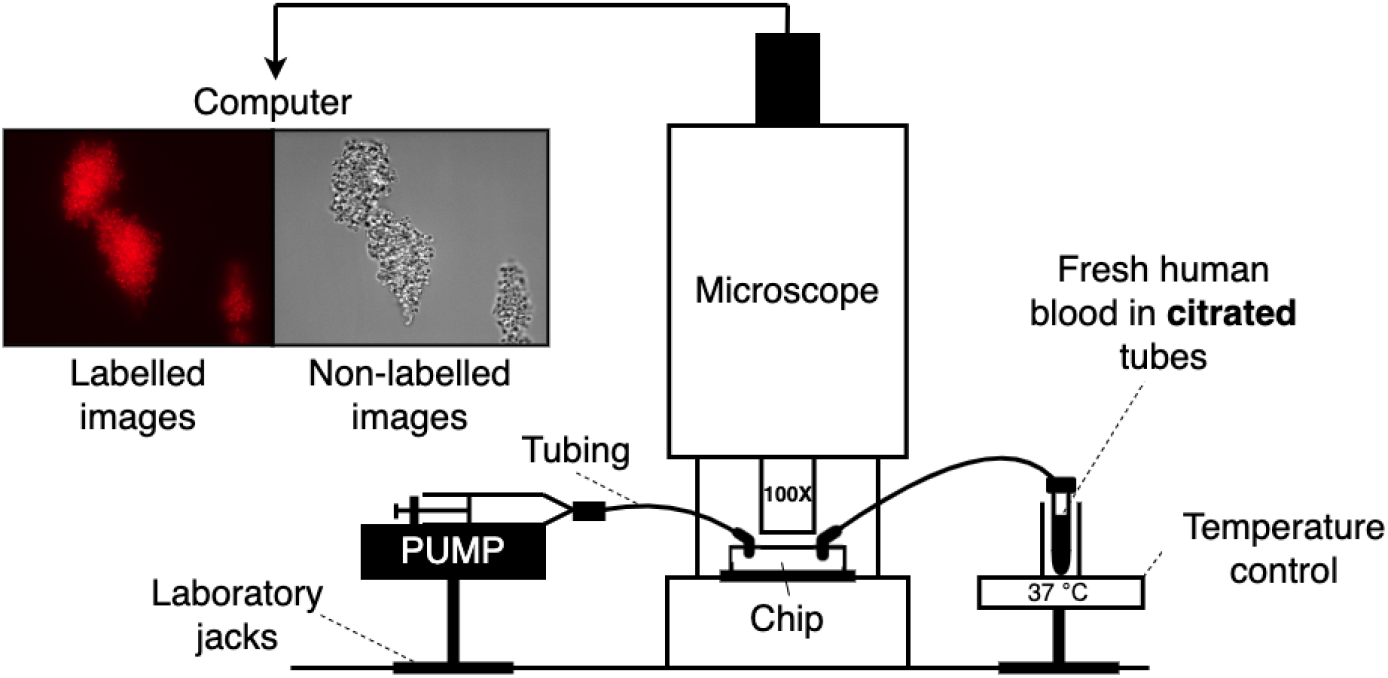
The schematic of the experiment set-up. The width and the height of the chamber is 1 *mm* and 100 *µm*, respectively. The dimension of the recorded experimental domain is 125 *µm ×* 100 *µm ×* 100 *µm*.

### Image data processing

A schematic workflow of the methodology employed in this work to transfer the experimental image data to the computational model is provided in Fig 2.

**Fig 2.**
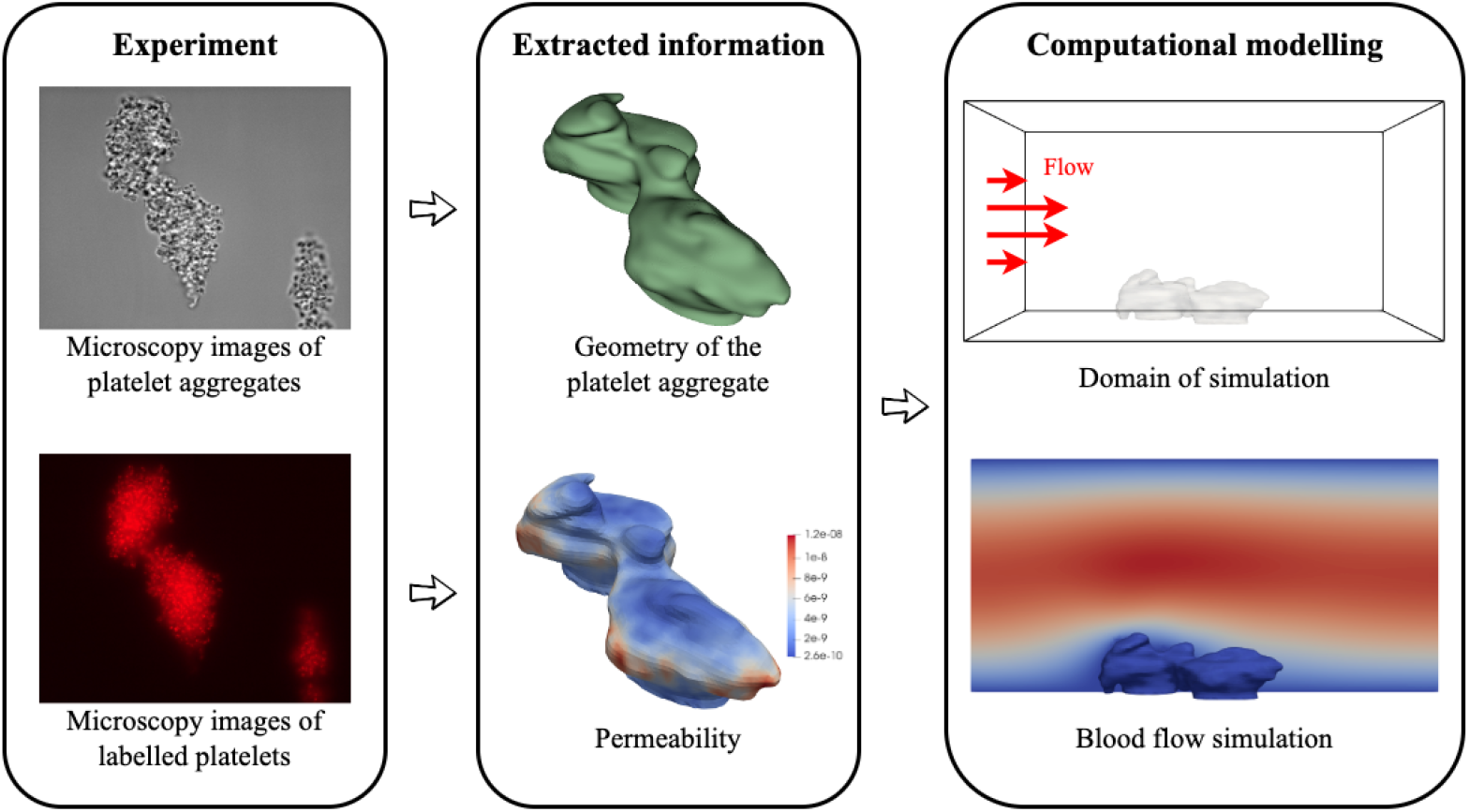
Flow diagram for the image-based modelling methodology implemented in this work. *Experiment* : microscopy images of platelet aggregates with labelled and non-labelled platelets caught by differential interference contrast microscopy with 100x magnification. *Extracted information*: reconstructed platelet aggregate geometry and permeability distribution over the aggregate. *Computational modelling* : the simulated domain for blood flow.

The non-labelled images were segmented with 3D Slicer [33] manually and stacked together over the z direction. As a final step to reconstruct the surface of the platelet aggregate, interpolation was applied to fill between the image slices. Smoothing was applied to further improve the surface quality of the platelet aggregate meshes.

However, the surface mesh may still contain low quality mesh elements, therefore it was remeshed with MeshLab [34] using the uniform resampling filter. After proceeding with the surface mesh, a corresponding volume mesh was created in Gmsh [35] for use in the numerical simulations.

The intensity of the fluorescence in the labelled image data is proportional to the local density of the platelet aggregate, and can be further used to infer the porosity of the aggregates. Therefore density measurements have been made to approximate the local density of aggregates. This was done by manually counting and the details of it are described in S1 File. Fig 3 shows the results of this measurement which provide us with a mapping between the fluorescence intensity and density of the entire aggregates. The permeability of the aggregates was subsequently inferred using the Kozeny-Carman equation. Further details on this are presented in the next subsection.

**Fig 3.**
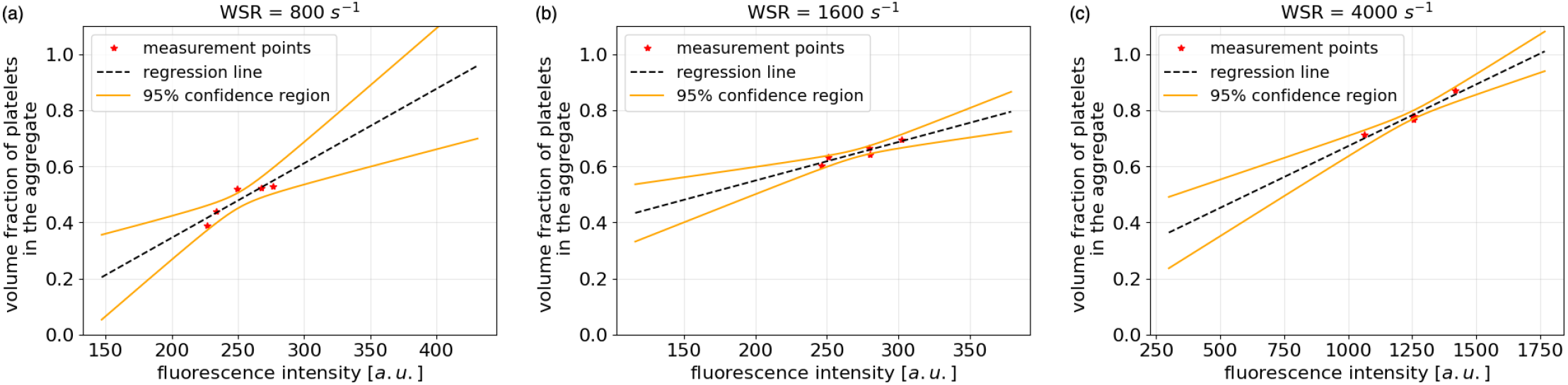
Fluorescence intensity (arbitrary units) - density relation inside platelet aggregates. (a) 800 *s*^−1^ WSR. Slope = 0.0027. Intercept = -0.19. (b) 1600 *s*^−1^ WSR. Slope = 0.0014. Intercept = 0.28. (c) 4000 *s*^−1^ WSR. Slope = 0.0004. Intercept = -0.23. The unit in each subfigure is different due to the influence of external environment such as exposure level and labelling time.

To simulate the blood flow around the platelet aggregates, a rectangular domain was created and uniformly meshed to represent a part of the channel of the chip in the experiments, as shown in Fig 4. The platelet aggregates formed on the collagen coated bottom plate, approximately in the middle of the channel. The intensity data of platelet aggregates were interpolated from the platelet aggregate volume mesh to the volume mesh of the flow domain. This interpolation between two meshes allows us to assign the corresponding fluorescence intensity value of the aggregate to a specific location in the simulated domain. Finally, this volume mesh of the flow domain was then used as the spatial discretization of the finite element method to simulate the blood flow.

**Fig 4.**
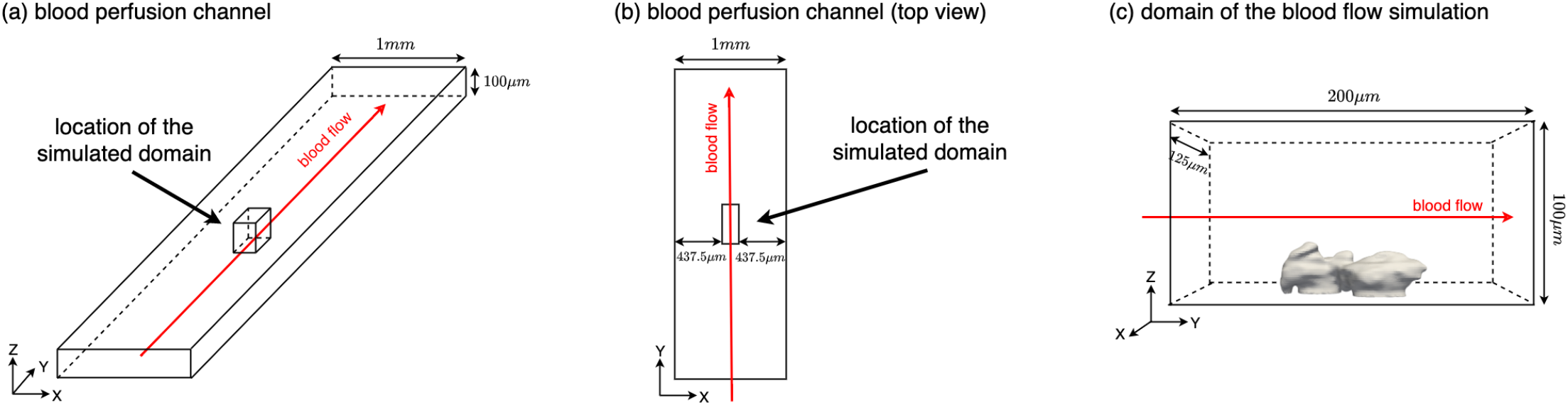
Schematic diagram of the simulation domain. (a) Blood perfusion channel and the location of the simulated domain. (b) Top view of the blood perfusion channel. (c) Domain of the blood flow simulation including the position of the platelet aggregate.

### Computational blood flow model

Blood flow was modelled as an incompressible Newtonian fluid governed by the Navier-Stokes equations, and the influence of the porous clot on the flow was introduced using a Darcy term:

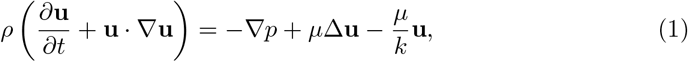

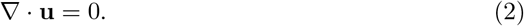

Here, **u** is the velocity of the fluid, *p* is the pressure, *ρ* is the fluid density, and *µ* is the dynamic viscosity. The last term on the right-hand side accounts for the momentum exchange between the fluid and the solid phases of the porous medium. It is defined by Darcy’s law [36, 37], which describes flow through a porous medium. In this term, *k* is the permeability of the platelet aggregates. Permeability is a function of the volume fraction of pore size and fiber or cell/cell aggregate size [21] and was estimated using the Kozeny-Carman equation

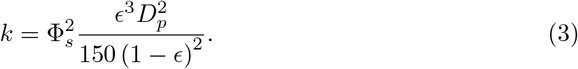

Here Φ_*s*_, *ϵ* and *D*_*p*_ are the sphericity of the platelets, the porosity of the aggregates and the platelet diameter, respectively.

The complete list of parameter values used in the model and their literature sources are shown in Table 1. According to the inlet flow settings in experiments, the Reynolds numbers for 800 *s*^−1^, 1600 *s*^−1^ and 4000 *s*^−1^ WSRs is 0.67, 1.37 and 3.33, respectively. We expect that the flow velocity in the platelet aggregates are significantly smaller than the velocity in the channel due to the porous structure, which fits the Stokes flow assumption of the Kozeny-Carman equation. The time-step size was chosen to satisfy the Courant–Friedrichs–Lewy condition. A mesh convergence study was carried out to select the appropriate mesh resolution for the simulation. Considering the computational cost and the convergence of the simulation, a mesh with 3.0 million tetrahedral elements was chosen. The average element size of the mesh is 0.837 *µm*^3^, which is significantly smaller than the average size of a platelet (7.2−11.7 *f L*) [38]. As shown in Fig 4, the blood flowed through the domain from left to right (along the positive y direction). Since the blood flow domain was considered as a small part of the channel in the x direction, constant and parabolic velocity profiles were prescribed in the x and z directions respectively at the inlet (for the coordinate system see Fig 4). A constant-pressure condition was prescribed at the outlet. No-slip boundary conditions were imposed at the top and bottom walls.

**Table 1.** Parameter used in simulations.

| Notation | Description | Value | Unit | Reference |
| --- | --- | --- | --- | --- |
| $\rho$ | density of fluid | $1.025 \times 10^{-3}$ | $g\ mm^{-3}$ | [39] |
| $\mu$ | fluid viscosity | $3 \times 10^{-3}$ | $g\ mm^{-1}\ s^{-1}$ | [32, 40] |
| $\Phi_s$ | sphericity of activated platelet | 0.71 | — | [41] |
| $D_p$ | platelet diameter | $2 \times 10^{-3}$ | $mm$ | [42] |
| $dt$ | time-step size | $10^{-5}$ | $s$ | — |
| — | average element size | 0.837 | $\mu m^3$ | — |

The flow field was solved using the finite element method (FEM) [43] and implemented in FreeFEM [44]. An iterative solver based on flexible generalized minimal residual (FGMRES) method [45] was used and implemented via PETSc [46]. This work was carried out on the Dutch national supercomputer Snelllius (SURF, Netherlands).

Each simulation was performed on AMD Rome 7H12 CPU × 2 and parallelized on 128 cores. The corresponding code of the simulation is available at https://github.com/UvaCsl/AggregateFlowSimu/releases/tag/v1.0.0.

## Results

The distribution of the fluorescence intensity obtained from the experimental images under three WSRs is presented in Fig 5 (a)-(c). With the increase of the shear rate, the ratio of fluorescence intensity clustering at the tail of the distribution increases, which means more parts in the aggregates have a higher density. Note that the values of this fluorescence intensity are in arbitrary units and not comparable under different scenarios due to the fact that they are influenced by other factors such as initial labeling concentration and light intensity. By computing the corresponding aggregate density from the fluorescence intensity via regression, the average density and the distribution of this density inside the aggregates are demonstrated in Fig 5(d) and Fig 5(e). Subsequently, Fig 5(f) shows the distribution of the permeability. The denser the aggregate is, the lower the permeability.

**Fig 5.**
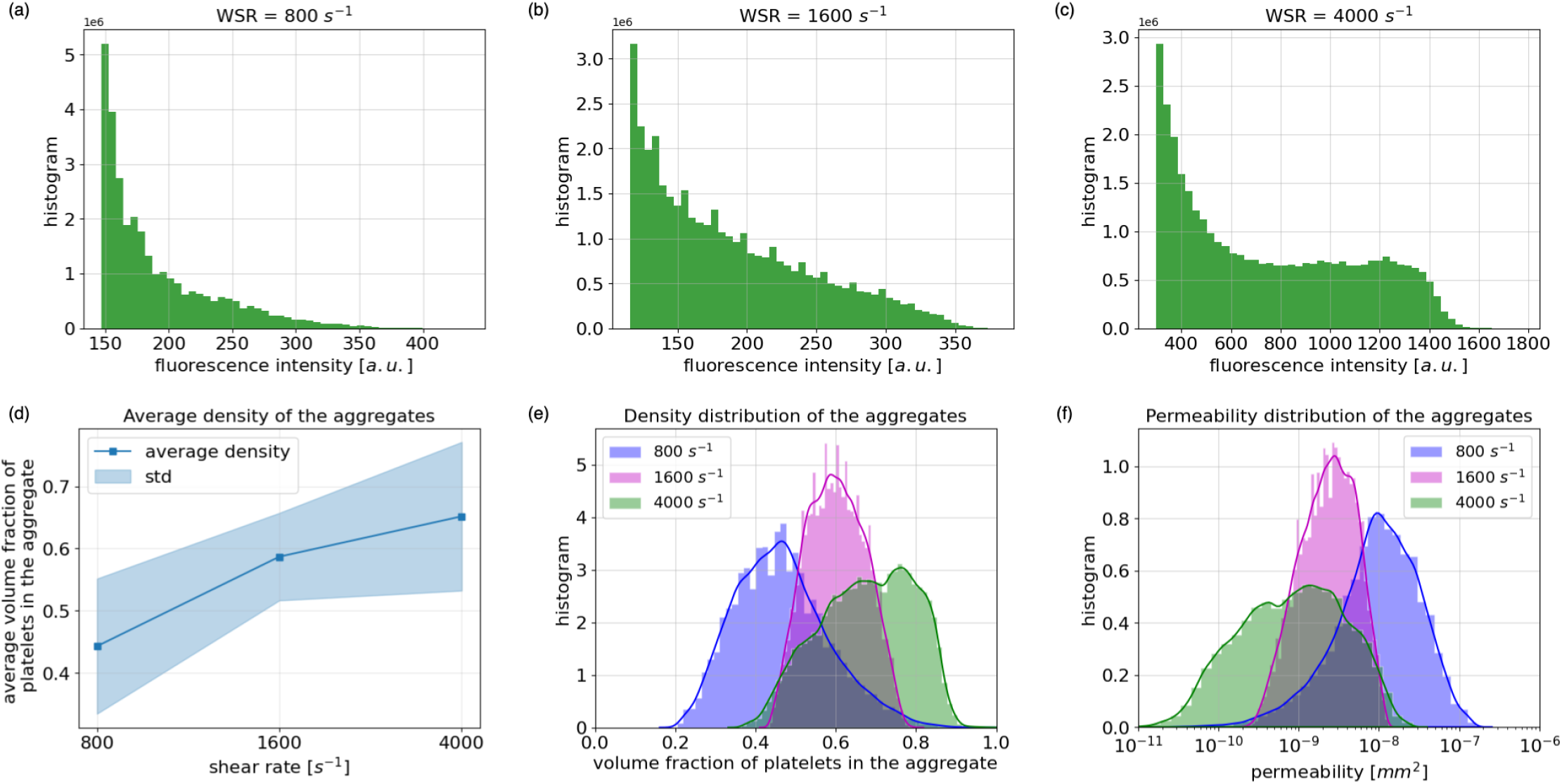
Distribution of fluorescence intensity, distribution of platelet aggregates density and permeability under 800 *s*^−1^, 1600 *s*^−1^ and 4000 *s*^−1^ WSRs. (a)-(c) Distributions of fluorescence intensity inside the platelet aggregates. (d) Average volume fraction of platelets of the platelet aggregates. (e) Distribution of volume fraction of platelets inside the platelet aggregates. (f) Distribution of permeability inside the platelet aggregates in log-scale.

To further study how fluorescence intensity, porosity and permeability distribute inside the platelet aggregate, the corresponding results on the cross-sections of the aggregate formed under 1600 *s*^−1^ WSR are shown in Fig 6. In the core of the aggregate, the intensity is higher than that in the shell. Correspondingly, the platelets in the core are positioned denser, which leads to much lower permeability in this part. This non-homogeneity results in complex blood flow behavior inside the platelet aggregates.

**Fig 6.**
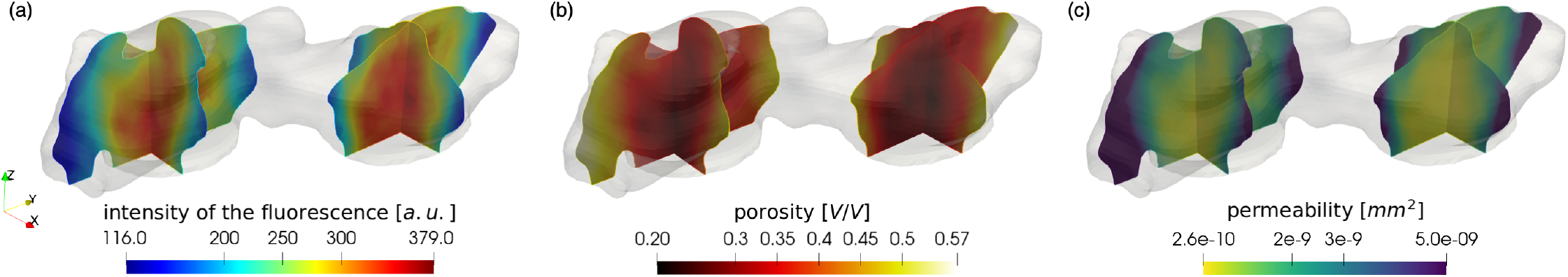
Intensity of the fluorescence, corresponding porosity and the permeability of the platelet aggregate under 1600 *s*^−1^ WSR. (a) Intensity of the fluorescence obtained from the experimental data. (b) Corresponding porosity of the platelet aggregate. (c) Permeability of the platelet aggregate on the cross-sections of the aggregate obtained from the Kozeny-Carman equation. The results inside the platelet aggregates formed under 800 *s*^−1^ and 4000 *s*^−1^ WSRs are demonstrated in S1 Fig.

## Flow within and around the platelet aggregate

The steady flow pattern on a cross-section under 1600 *s*^−1^ WSR is shown in Fig 7(a). Although the platelet aggregate is considered as a porous medium, the permeability of the aggregates is extremely small. Consequently, most of the blood flows over the aggregate and only a small amount of the blood permeates the platelet aggregate.

**Fig 7.**
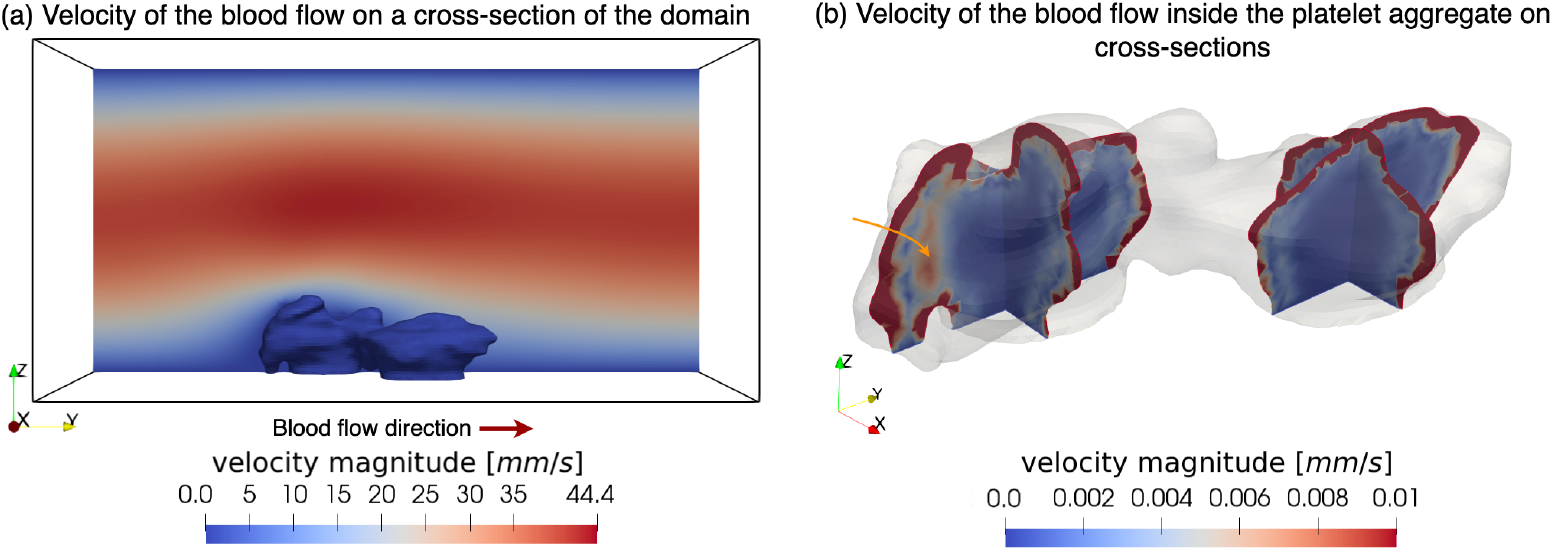
Flow field at WSR of 1600 *s*^−1^. (a) The velocity field of the blood flow on a cross-section of the flow domain. The red arrow indicates the direction of blood flow. (b) The velocity field of the blood flow inside the platelet aggregate on the cross-sections. The orange arrow points out the high flow velocity area between the shell and the core.

Fig 7(b) shows the corresponding blood flow velocity inside the platelet aggregates on cross-sections. Although in the shell with relatively high permeability, the blood flow velocity is significantly lower than the extra-thrombus flow, it is still considerably higher than the velocity in the core with lower permeability. However, in the transition from the shell to the core, the magnitude of the velocity is greater than that of some regions of the shell. After blood enters the core, its velocity becomes extremely low. The minimum velocity in the interior core reaches 10^−11^ *mm/s*, which means barely any blood flow.

### Stress analysis of the blood flow

Since the platelet aggregate is considered as a porous medium, the momentum of the flow exerts a kinetic force inside the aggregate, which can lead to partial or complete aggregate embolism [32]. These forces indicate the interaction between blood flow and platelet aggregates under the assumption that the deformation of the aggregate is insignificant, and induce stresses in the aggregate structure. However, these are not discussed in the current work. This kinetic force was calculated by Darcy’s law, 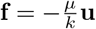, and the result is shown in Fig 8(a). The highest forces appear on the top outer layer of the platelet aggregate, due to the relatively high fluid velocity over these parts. Also, similar to the blood velocity in the transition from the shell to the core, there exists an area with higher kinetic force compared to the force in the shell and core.

**Fig 8.**
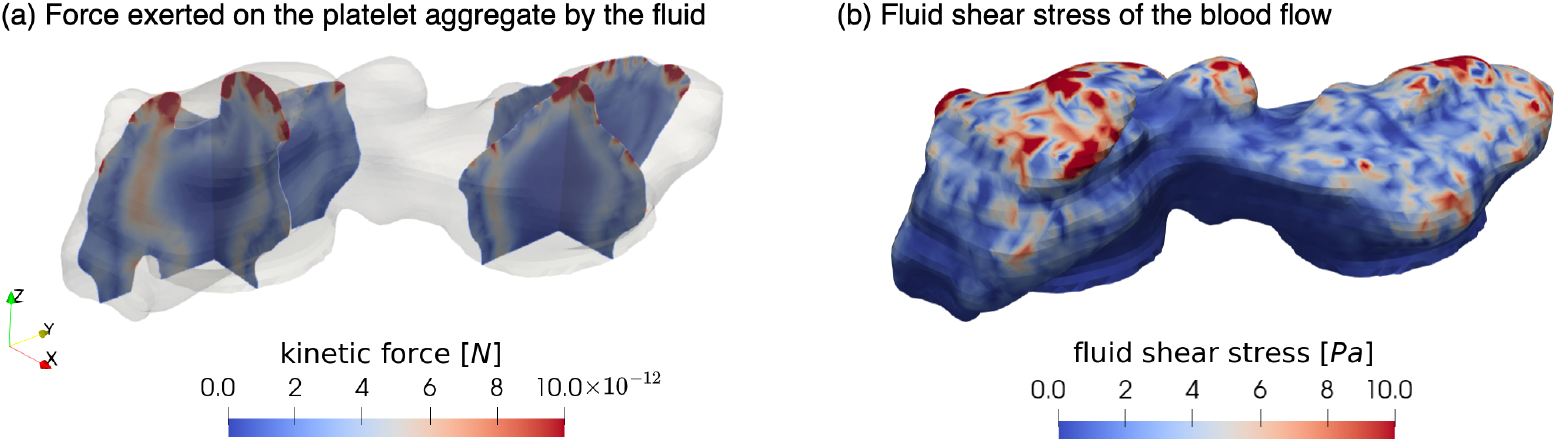
Stress analysis of the blood flow and the platelet aggregate under 1600 *s*^−1^ WSR. (a) The kinetic force exerted on the platelet aggregate. (b) The fluid shear stress on the surface of the platelet aggregate. The simulation results inside the platelet aggregates formed under 800 *s*^−1^ and 4000 *s*^−1^ WSRs are shown in S2 Fig.

Furthermore, the local shear stress has a significant influence on platelet activation, aggregation and adhesion [47]. It has been shown in [48] and [49] that high levels of shear stress could initiate platelet aggregation. Fig 8(b) demonstrates the fluid shear stress on the surface of such platelet aggregate under 1600 *s*^−1^ WSR condition. These fluid-induced shear stress magnitude *σ* is estimated by:

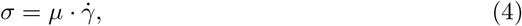

where 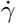 is the shear rate magnitude obtained by:

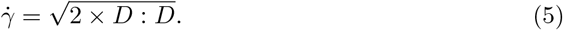

Here, *D* represents the strain rate tensor which is defined as 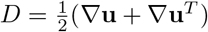. As expected, shear stress at the top part of the aggregate is several-fold higher than at the bottom. This is because the aggregate protrudes into the center of the channel and is exposed to a higher blood flow velocity. This result agrees with the shear stress distribution reported in [28] and [32].

### Advection versus diffusion

Biochemical processes involving different agonists such as adenosine-5^*′*^-diphosphate (ADP), require a certain amount of time to take place. In the advection-dominated region, if such a process takes more time than the characteristic time of agonist advection (i.e. the average time of the agonist to be convected over a distance of platelet), this process is hindered since the flow velocity is high enough to wash away the agonists before any reaction takes place. Therefore, identifying the advection-dominated region is critical for understanding the dynamics of the aggregation process. The advection- and diffusion-dominated regions of inner chemicals were evaluated by Péclet number, a ratio of advection and diffusion time:

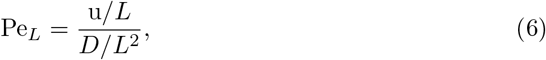

where u denotes the magnitude of local flow velocity, *D* is the mass diffusion coefficient and *L* denotes the characteristic length. In our simulation, the characteristic length is defined as the platelet diameter, therefore the advection time in the denominator of Pe_*L*_ indicates the time required to move the distance of a platelet diameter. When the Péclet number equals one, advection and diffusion contribute to mass transport equally. If Pe_*L*_ *>* 1, the diffusion time of the chemical is longer than the advection time in such area, which means the motion of the chemical is advection-dominated. Otherwise, it is diffusion-dominated.

Three agonists, Calcium (Ca^2+^), ADP and Factor X, were evaluated in this work (see Table 2). Ca^2+^ contributes to multiple stages of cellular activation in platelets [50]. It has the lowest molecular weight of these three chemicals but the highest diffusion coefficient. The second one is ADP, an agonist that plays an important role in platelet activation. Finally, Factor X, an enzyme in the coagulation cascade that can increase the thrombosis propensity [51]. It has the highest molecular weight and the lowest diffusion coefficient among the three agonists. The results demonstrate that the advection-dominated volume in the platelet aggregate increases with the decrease of the diffusion velocities of agonists. Furthermore, Fig 9 visualises the distribution of the advection- and diffusion-dominated areas under the same flow condition. It can be observed that for the highly diffusive agonists only minuscule parts of the platelet aggregate are advection-dominated, even in the shell region. However for Factor X, advection takes over the transport at almost the entire outer layer of the aggregate. In conclusion, with the decrease of the diffusion coefficient, advection gradually dominates the transport of chemicals in the shell region of the platelet aggregate. This aligns well with previous findings predicting such behaviour within the core-shell structure of platelet aggregates [13, 17].

**Table 2.**
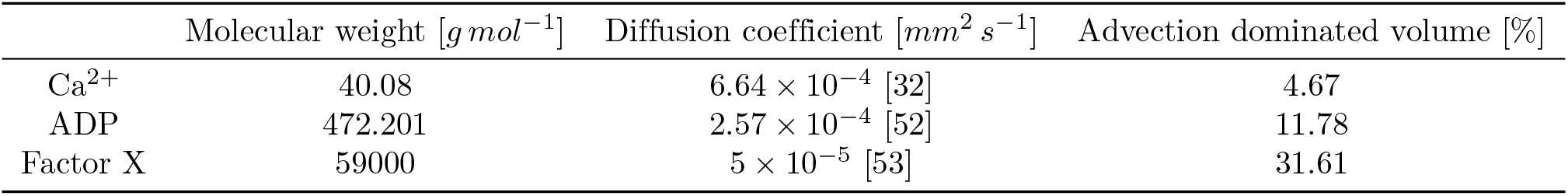
Intrathrombus transport simulation for coagulation factors with different molecular weights at WSR of 1600 *s*^−1^.

| | Molecular weight [$g\ mol^{-1}$] | Diffusion coefficient [$mm^2\ s^{-1}$] | Advection dominated volume [%] |
| --- | --- | --- | --- |
| $Ca^{2+}$ | 40.08 | $6.64 \times 10^{-4}$ [32] | 4.67 |
| ADP | 472.201 | $2.57 \times 10^{-4}$ [52] | 11.78 |
| Factor X | 59000 | $5 \times 10^{-5}$ [53] | 31.61 |

**Fig 9.**
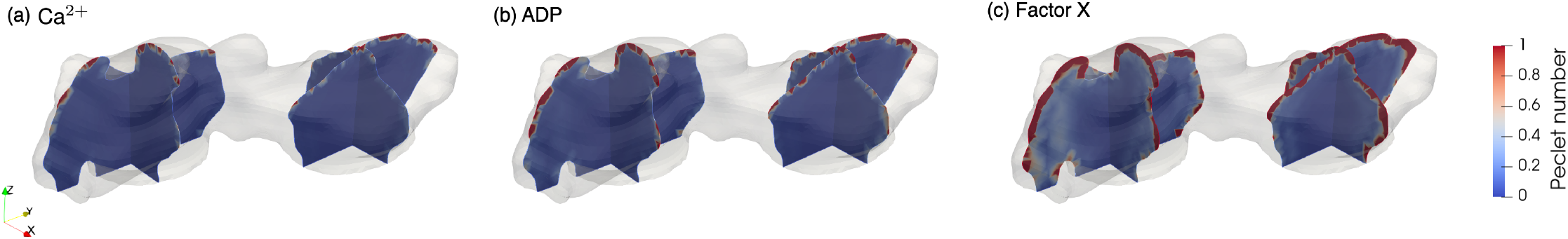
Advection-diffusion balance. Advection-diffusion balance of (a) Ca^2+^, (b) ADP and (c) Factor X on cross-sections inside the platelet aggregate under 1600 *s*^−1^ WSR. The upper end of the color scale is set to 1. Therefore, areas with red color correspond to advection-dominated regions, while colors towards the lower end of the scale denote diffusion-dominated regions.

### Effects of increasing WSR

Under all three shear rate conditions, the average intrathrombus velocity is at least two orders of magnitude lower than the average extra-thrombus velocity, as shown in Table 3. This is comparable to the previously reported results on the microscale thrombus aggregate simulation [16, 54]. Furthermore, the increase in inlet blood velocity leads to higher average intrathrombus velocity and stronger kinetic forces on platelet aggregates.

**Table 3.** Comparison of simulation results under various WSRs.

| Wall shear rate [$\text{s}^{-1}$] | | 800 | 1600 | 4000 |
| --- | --- | --- | --- | --- |
| Intrathrombus velocity [$\text{mm s}^{-1}$] | | $0.05 \pm 0.09$ | $0.06 \pm 0.15$ | $0.13 \pm 0.34$ |
| Extra-thrombus velocity [$\text{mm s}^{-1}$] | | $13.78 \pm 6.28$ | $27.50 \pm 12.79$ | $69.06 \pm 32.18$ |
| Kinetic force [$N$] ($\times 10^{-12}$) | | $1.36 \pm 2.07$ | $4.07 \pm 5.34$ | $11.20 \pm 19.34$ |
| Average Péclet number | $\text{Ca}^{2+}$ | $0.16 \pm 0.27$ | $0.18 \pm 0.44$ | $0.38 \pm 1.02$ |
| | ADP | $0.40 \pm 0.69$ | $0.45 \pm 1.13$ | $0.99 \pm 3.64$ |
| | Factor X | $2.08 \pm 3.56$ | $2.33 \pm 5.80$ | $5.11 \pm 13.59$ |
| Advection-dominated volume [%] | $\text{Ca}^{2+}$ | 1.97 | 4.67 | 10.90 |
|  | ADP | 14.46 | 11.78 | 18.82 |
|  | Factor X | 37.82 | 31.61 | 33.83 |

Average intrathrombus velocity, average extra-thrombus velocity, average kinetic force, average Péclet number inside the aggregates and the advection-dominated volume of $\text{Ca}^{2+}$ , ADP and Factor X inside the platelet aggregates under three WSRs.

In order to explore how the inlet WSRs affect the solute transport, the average Péclet number and advection-dominated volume inside the platelet aggregates are also computed. The result indicates that the average Péclet number inside the aggregates has a positive correlation with WSRs, while the relation between advection-dominated volume and WSRs is not clear.

To further understand the influence of the blood flow velocity on the platelet aggregate formation, the shapes of the platelet aggregates formed under three shear rates are shown in Fig 10. It can be observed that the platelet aggregate formed under a high shear rate condition (4000 *s*^−1^) has a distinctly different shape. It is the tallest out of the three, while the shape of the aggregate formed under low shear rates is the flattest.

**Fig 10.**
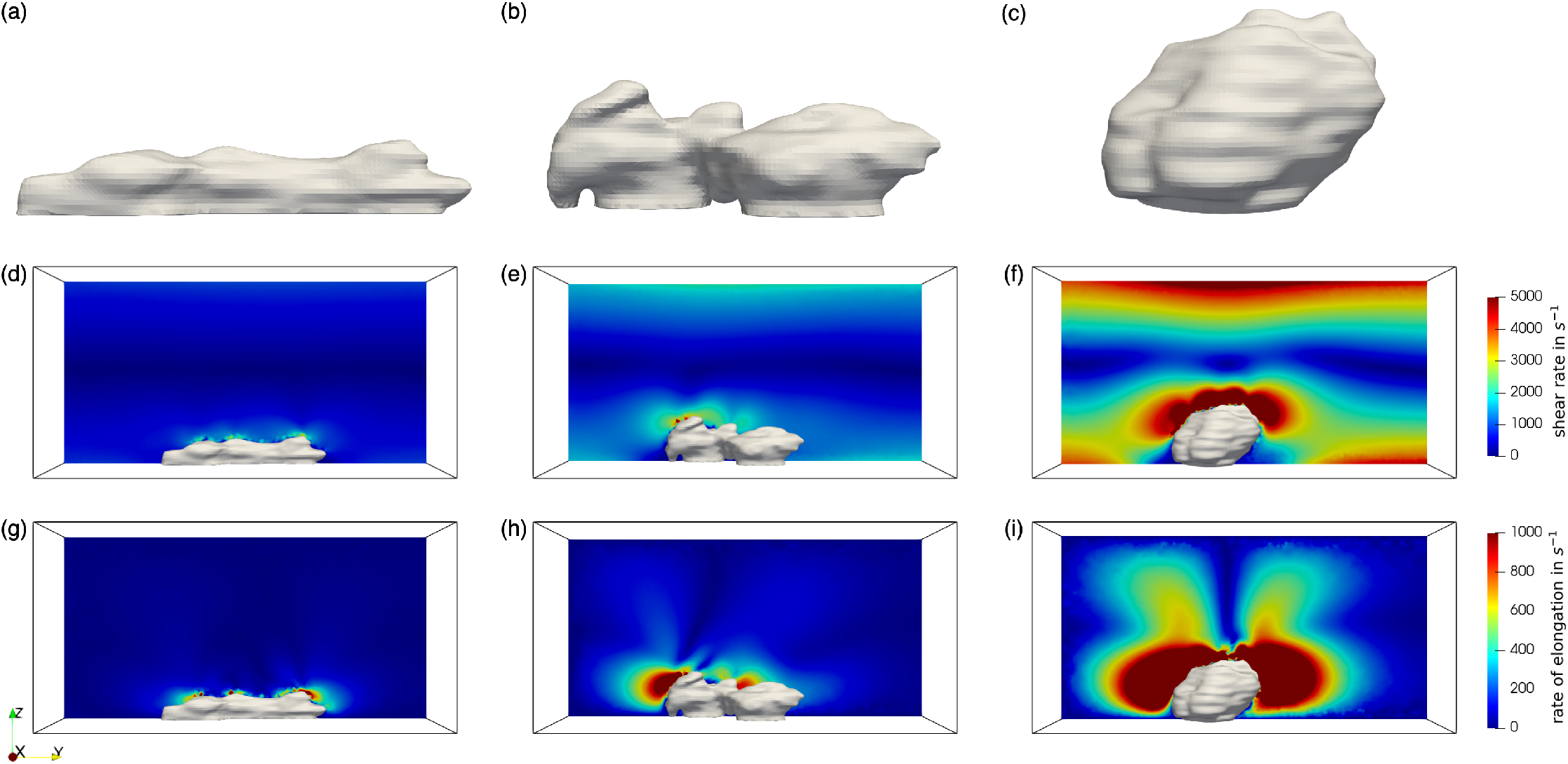
Platelet aggregates geometries, shear rates and the rate of elongation. (a)-(c) Geometries of the platelet aggregates under 800 *s*^−1^, 1600 *s*^−1^ and 4000 *s*^−1^ WSRs flow condition. (d)-(f) Cross-sectional shear rate profile under 800 *s*^−1^, 1600 *s*^−1^ and 4000 *s*^−1^ WSRs flow condition. (g)-(i) Cross-sectional elongation rate profile under 800 *s*^−1^, 1600 *s*^−1^ and 4000 *s*^−1^ WSRs flow condition.

Platelet adhesion has been demonstrated to be strongly influenced by the mechanosensitivity of VWF [55–58]. This protein uncoils and activates only under certain flow conditions [59]. Previous studies have shown that VWF will unfold when the shear rate or the rate of elongation exceeds a threshold [59, 60]. Therefore, high shear rate and elongation rate facilitate the binding of platelets and can lead to platelet activation [61]. The appearing shear rate and elongational rate in the three cases are investigated in Fig 10. The shear rate magnitude *γ*? is computed based on the corresponding flow rates:

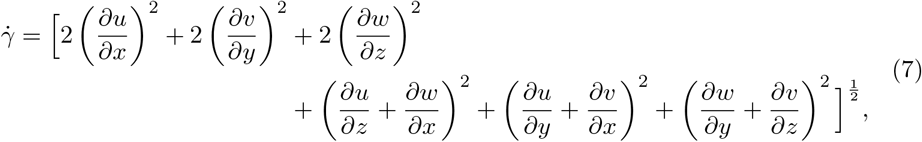

where *u, v* and *w* are the flow velocity in the x-, y- and z-directions, respectively. The rate of the uni-axial elongation is defined as the magnitude of the diagonal elements of the rate of the flow strain tensor [62, 63]:

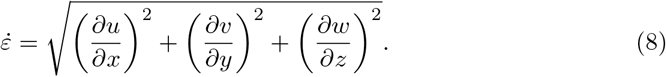

According to [59, 60, 64], the threshold of elongation rate for VWF to unfold is much smaller than that of the shear rate: when the shear rates reach 5000*s*^−1^ or elongation flow strain rates are higher than 1000 *s*^−1^, the compact conformation of VWF will change [60, 65]. Hence, to observe the region where VWF unfolds in our cases, the color bar maxima for results of the shear rate and the elongational flow are set to coincide with the thresholds reported by these studies, namely 5000 *s*^−1^ and 1000 *s*^−1^, respectively. The figure indicates that under a high shear rate condition (4000 *s*^−1^), the region where the shear rate or the elongation rate exceeds the threshold, is considerably larger than the region in the case of low shear rate flow conditions. This leads to a higher potential for platelets to aggregate and adhere under high shear rates.

### Comparison of different ranges of porosity

In this work, the porosity ranges are inferred from the local density of the platelets. The measurement requires one to count the number of platelets manually in multiple regions of the aggregate, which causes uncertainty (for further details see S1 File). Furthermore, aggregates formed under different conditions might display different porosities.

Therefore, a series of simulations based on different porosity ranges were performed to quantify the possible influence of potential porosity ranges on the quantities of interest. Different porosity values and different ranges of porosity were chosen and considered to have a linear relation with fluorescence intensity. Fig 11 visualises the changes in average blood flow velocity and advection-dominated volume inside the aggregates in terms of a fixed range of porosity under three WSRs. With the same porosity settings, both the average velocity and the advection-dominated volume are higher under a higher WSR, which is expected. Simultaneously, the increase of the porosity, which means more permeable aggregates, leads to an increase of not only the average velocity but also more advection-dominated areas inside the aggregates. Moreover, the blood flow behaviour under various ranges of porosity was also investigated and the results are shown in Fig 12. With a wider porosity setting, both the average velocity within the aggregates and the effect of advection on chemical transport increase. Overall, these changes in the porosity lead to quantitative changes in the flow field, however, the observed trends remain unaffected, implying the wider generality of the results.

**Fig 11.**
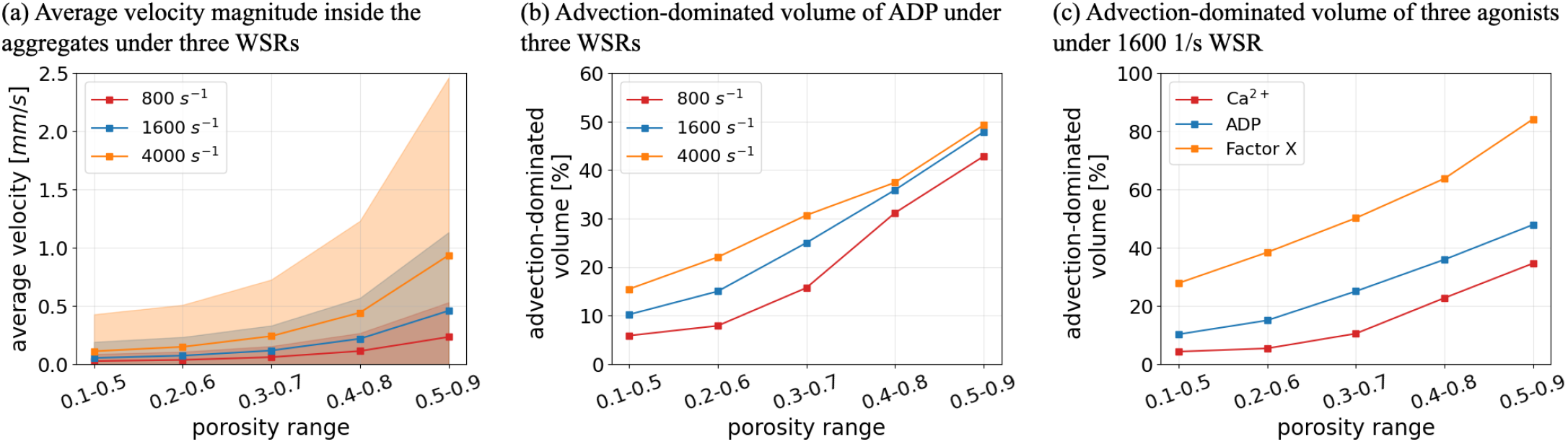
Comparison of intrathrombus condition for different porosity values. (a) Average blood flow velocity inside the platelet aggregates under various shear flow conditions. (b) Advection-dominated volume of ADP inside the platelet aggregates under various shear flow conditions. (c) Advection-dominated volume of Ca^2+^, ADP and Factor X inside the platelet aggregates under WSR of 1600 *s*^−1^.

**Fig 12.**
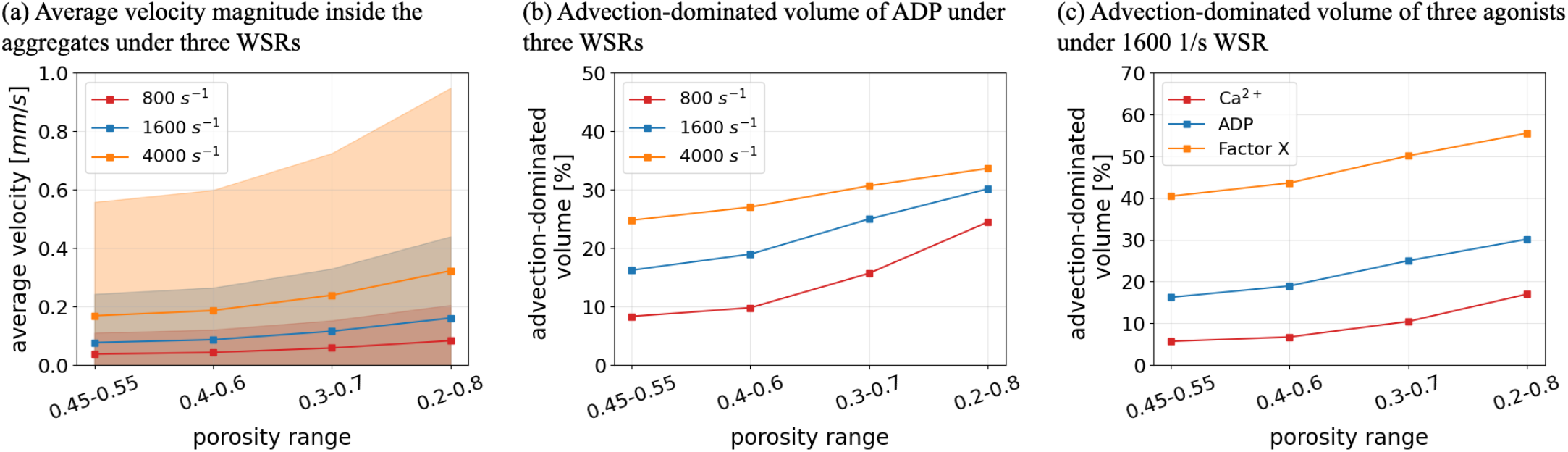
Comparison of intrathrombus condition for different porosity ranges. (a) Average blood flow velocity inside the platelet aggregates under various shear flow conditions. (b) Advection-dominated volume of ADP inside the platelet aggregates under various shear flow conditions. (c) Advection-dominated volume of Ca^2+^, ADP and Factor X inside the platelet aggregates under WSR of 1600 *s*^−1^.

### Limitations

The proposed image-based computational model is based on several assumptions and has some limitations. First, the porosity of the aggregates was inferred from the fluorescence intensity of the images. However, the exposure time and exposure level in the experiments can have an impact on the fluorescence intensity. How the fluorescence intensity changes with the exposure time and level should also be investigated in more detail to provide a more precise estimation of the platelet density. Second, the porosity of the platelet aggregates is approximated by the ratio of the area occupied by platelets on the images. This process requires manually counting the number of platelets, which involves uncertainty and contributes to the uncertainty in the model outputs. However, the evaluation of interobserver variability was not performed. Therefore, a comparison study based on different porosity settings was performed, and demonstrated how the potential uncertainty in porosity ranges would affect quantities of interest, such as average velocities and advection-dominated volume of agonists under three WSRs. No qualitative changes have been observed due to the variety of porosity ranges.

Furthermore, the volume fraction result shown in Fig 6 follows our expectation on platelet aggregates formed under different shear rates, i.e. the aggregates tend to be denser when formed under higher shear rates. Moreover, in the computational model of blood flow, the viscosity in the lumen was assumed to be constant. However, in actual flow, red blood cells in the microchannel migrate toward the center of the lumen resulting in a red blood cell-free layer close to the vessel wall [66, 67]. Such a phenomenon contributes to heterogeneity of viscosity in the lumen and therefore influences the flow resistance and biological transport [68]. Also, the platelets and red blood cells cannot penetrate platelet aggregates, which means the fluid inside the aggregates is pure plasma. Its viscosity is smaller than whole blood [69, 70]. A simulation with plasma viscosity has been carried out for comparison. The result (S3 Fig) shows that there is no significant influence to the flow dynamics inside the aggregates. Finally, the platelet aggregate pore wall presents barriers to the agonists, resulting in a smaller agonist diffusion coefficient, which contributes to a larger Péclet number. Since the knowledge of how large the effect of the porous on the diffusion of agonists is unknown, we used the diffusion coefficients that were measured in free flow.

## Discussion

In this work, two different microscopy image modalities were used to capture the morphology and microstructure of platelet aggregates formed under *in vitro* blood perfusion experiments. The information extracted from the images yields the geometry and local porosity of the aggregates, that are subsequently applied in the computational model. The image-based computational model enables an accurate prediction of intrathrombus flow under various flow conditions. A separation of the core and shell of the aggregates in terms of the platelet density can be observed in Fig 6. The core is composed of densely packed platelets, where the transport of agonists is diffusion-dominated, while the outer layer is a loose association of platelets.

Furthermore, the blood flow velocity inside platelet aggregates observed has a qualitative agreement with the result reported in [32]. However, a strip area with relatively high flow velocity was found at the boundary between the shell and the core. To understand what happens here, the magnitude of the blood flow velocity under three WSRs was investigated. As Fig 13 shows, this situation occurs under all three WSRs. This might be because the platelets in the core were excessively dense, causing the circumferential flow around the core and contributing to the increase of the blood flow velocity at the transition zone from shell to core. Correspondingly, the large kinetic forces in the area were generated by the resistance when blood flowed from the shell to the core, a much denser medium. This phenomenon may help to identify the boundary between the shell and the core.

**Fig 13.**
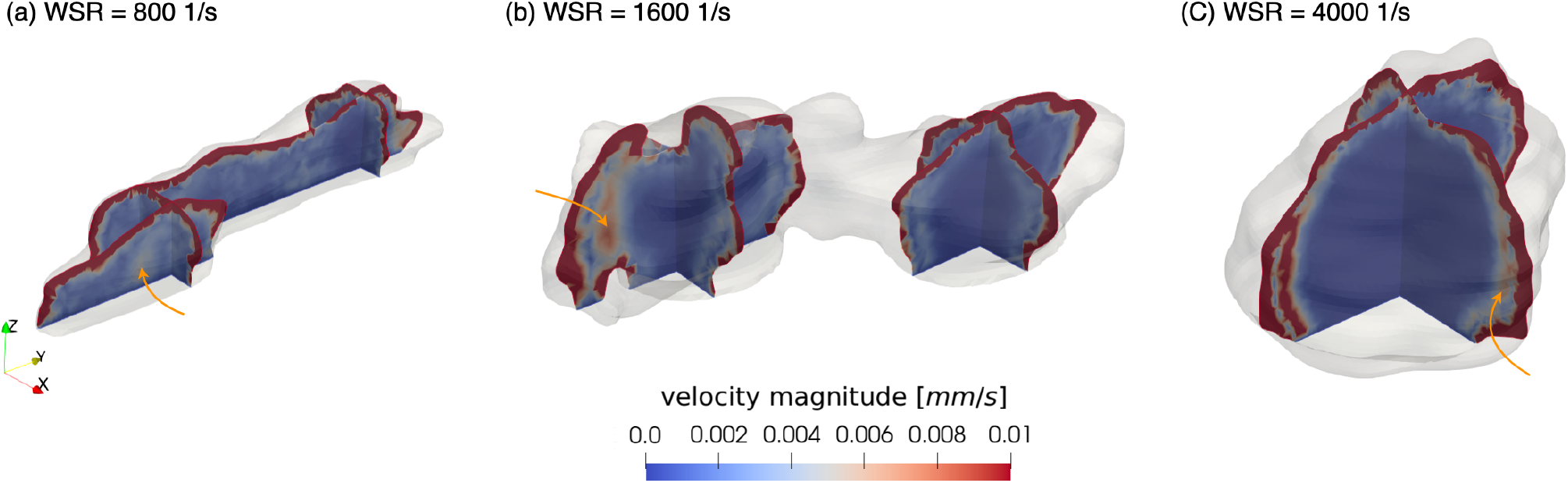
Velocity magnitude of the blood flow inside the platelet aggregates under three WSRs. (a) 800 *s*^−1^, (b) 1600 *s*^−1^ and (c) 4000 *s*^−1^. The arrows point out the high flow velocity area between the shell and the core.

In Table 3, no clear relation is found between advection-dominated volume and WSRs. This is mainly due to the density difference of the aggregates formed under three WSRs. As mentioned before, the platelet aggregate is denser under 4000 *s*^−1^ WSR, implying that there are more regions in the aggregate that large proteins, e.g. Factor X, cannot advect by the fluid. Therefore, this leads to a smaller advection-dominated volume for Factor X under 4000 *s*^−1^ WSR than that under 800 *s*^−1^ WSR. In contrast, for the lighter agonists such as Ca^2+^, the transport is not significantly hampered even if the aggregate becomes relatively denser in the core. As a result, advection dominates more area for the transport of lighter agonists inside the platelet aggregates with the increase of the blood shear rate.

As mentioned before, if the shear rate or the rate of elongation exceeds a threshold, the VWF will unfold and expose the A1 domain, hence supporting platelet attachment. As shown in Fig 10, under high shear rate flow conditions, the shear rate and the rate of elongation of most of the area around the aggregate exceed the threshold, which means the aggregates are exposed to a large amount of unfolded VWF. This might contribute to the easy attachment of VWF to the surface of aggregates, especially on the top of the aggregates. It further explains the phenomenon that the aggregates are prone to grow in parallel to the flow under a low shear rate while the ones formed under a high shear rate show an aggregation tendency perpendicular to the flow direction. The results of the shear rate and the rate of elongation might be further used to predict the aggregation process over time. However, this observation on clot geometries is based on a limited number of experiments. More experiments are required to show if it has statistical significance.

In this work, we considered the aggregates as porous media and mainly focused on the blood flow behaviour and its influence on the aggregates. However the aggregate might deform, contract, or even break away during this dynamic process. A model could be further developed to study the mechanical behaviour of the platelet aggregates. The kinetic force exerted on the platelet aggregates by the blood flow calculated from our model can be applied to this mechanical model. Combined with the experiments, the break-away dynamics of the aggregates and the mechanical stress forces under different environments can be further simulated and analyzed.

## Conclusion

In this work, we present an image-based computational blood flow model to simulate the blood flowing through the platelet aggregates which are considered as porous media. The shapes and the microstructures of the platelet aggregates are extracted from microscopy image data of platelet aggregates formed in the blood perfusion experiments. The fluorescence intensity of the platelet labelling is used to infer the porosity of the aggregates and the corresponding permeability is calculated via the Kozeny-Carman equation. The imaged-base model allows us to study the details of hemodynamic transport, such as the velocity, shear stress, and advection-diffusion of agonist transport inside and around the platelet aggregates. Relatively high blood flow velocity and forces were found at the transition from the shell to the core due to the compact distribution of platelets in the interior core. The large forces in the area could contribute to the activation of the platelets in the shell. Furthermore, the results demonstrate that both the flow shear rate and aggregate microstructure have a substantial impact on the transport of agonists. Finally, the shear rate and the rate of elongation flow have also been investigated and the findings imply that there is a strong correlation between these values and the shapes of growing platelet aggregates. Overall, the proposed computational model incorporates the internal microstructure of the aggregates and enables a more precise prediction of the hemodynamics in the platelet aggregates. Our work lays the foundation for predicting shear-dependent aggregation and deformation under different flow conditions, which could subsequently contribute to developing shear-selective anti-thrombotic or anti-platelet drugs.

## Supporting information

**S1 Fig. Intensity of the fluorescence, corresponding porosity and the permeability of the platelet aggregates under 800** *s*^−1^ **and 4000** *s*^−1^ **WSRs**. (a)-(c) Intensity of the fluorescence, corresponding porosity and permeability of the platelet aggregate on the cross-sections of the aggregate under 800 *s*^−1^ WSR. (d)-(f) Intensity of the fluorescence, corresponding porosity and permeability of the platelet aggregate on the cross-sections of the aggregate under 4000 *s*^−1^ WSR.

**S2 Fig. Stress analysis of the blood flow and the platelet aggregate under 800** *s*^−1^ **and 4000** *s*^−1^ **WSRs**. (a)-(b) The kinetic force exerted on the platelet aggregate and the fluid shear stress on the surface of the platelet aggregate under 800 *s*^−1^ WSR. (c)-(d) The kinetic force exerted on the platelet aggregate and the fluid shear stress on the surface of the platelet aggregate under 4000 *s*^−1^ WSR.

**S3 Fig. Flow field at WSR of 1600** *s*^−1^ **with plasma viscosity**. (a) The velocity field of the blood flow on a cross-section of the flow domain. (b) The velocity field of the blood flow inside the platelet aggregate on the cross-sections.

**S1 File. Platelet density measurement**

## References

1. Ni H, Freedman J. Platelets in hemostasis and thrombosis: role of integrins and their ligands. Transfusion and Apheresis Science. 2003;28(3):257–264. doi:https://doi.org/10.1016/S1473-0502(03)00044-2.

2. Nesbitt WS, Westein E, Tovar-Lopez FJ, Tolouei E, Mitchell A, Fu J, et al. A shear gradient–dependent platelet aggregation mechanism drives thrombus formation. Nature Medicine. 2009;15(6):665–673. doi:10.1038/nm.1955.

3. Hou Y, Carrim N, Wang Y, Gallant RC, Marshall A, Ni H. Platelets in hemostasis and thrombosis: Novel mechanisms of fibrinogen-independent platelet aggregation and fibronectinmediated protein wave of hemostasis. The Journal of Biomedical Research. 2015;29(jbr150602):437. doi:10.7555/JBR.29.20150121.

4. Jackson SP. The growing complexity of platelet aggregation. Blood. 2007;109(12):5087–5095. doi:https://doi.org/10.1182/blood-2006-12-027698.

5. Goto S, Ikeda Y, Saldívar E, Ruggeri ZM. Distinct mechanisms of platelet aggregation as a consequence of different shearing flow conditions. The Journal of Clinical Investigation. 1998;101(2):479–486. doi:10.1172/JCI973.

6. Savage B, Cattaneo M, Ruggeri Z. Mechanisms of platelet aggregation. Current opinion in hematology. 2001;8(5):270–276. doi:10.1097/00062752-200109000-00002.

7. Fogelson AL, Tania N. Coagulation under flow: the influence of flow-mediated transport on the initiation and inhibition of coagulation. Pathophysiology of Haemostasis and Thrombosis. 2005;34(2-3):91–108. doi:10.1159/000089930.

8. Yazdani A, Li H, Humphrey JD, Karniadakis GE. A general shear-dependent model for thrombus formation. PLOS Computational Biology. 2017;13(1):1–24. doi:10.1371/journal.pcbi.1005291.

9. Tokarev A, Sirakov I, Panasenko G, Volpert V, Shnol E, Butylin A, et al. Continuous mathematical model of platelet thrombus formation in blood flow. Russian Journal of Numerical Analysis and Mathematical Modelling. 2012;27(2):191–212. doi:doi:10.1515/rnam-2012-0011.

10. Mody NA, King MR. Platelet adhesive dynamics. Part I: characterization of platelet hydrodynamic collisions and wall effects. Biophysical Journal. 2008;95(5):2539–2555. doi:https://doi.org/10.1529/biophysj.107.127670.

11. Mody NA, King MR. Platelet adhesive dynamics. Part II: high shear-induced transient aggregation via GPIbα-vWF-GPIbα bridging. Biophysical Journal. 2008;95(5):2556–2574. doi:https://doi.org/10.1529/biophysj.107.128520.

12. Storti F, van Kempen THS, van de Vosse FN. A continuum model for platelet plug formation and growth. International Journal for Numerical Methods in Biomedical Engineering. 2014;30(6):634–658. doi:https://doi.org/10.1002/cnm.2623.

13. Leiderman K, Fogelson AL. Grow with the flow: a spatial–temporal model of platelet deposition and blood coagulation under flow. Mathematical Medicine and Biology: A Journal of the IMA. 2010;28(1):47–84. doi:10.1093/imammb/dqq005.

14. Leiderman K, Fogelson AL. The influence of hindered transport on the development of platelet thrombi under flow. Bulletin of Mathematical Biology. 2013;75(8):1255–1283. doi:10.1007/s11538-012-9784-3.

15. Welsh JD, Stalker TJ, Voronov R, Muthard RW, Tomaiuolo M, Diamond SL, et al. A systems approach to hemostasis: 1. The interdependence of thrombus architecture and agonist movements in the gaps between platelets. Blood. 2014;124(11):1808–1815. doi:10.1182/blood-2014-01-550335.

16. Mirramezani M, Herbig BA, Stalker TJ, Nettey L, Cooper M, Weisel JW, et al. Platelet packing density is an independent regulator of the hemostatic response to injury. Journal of Thrombosis and Haemostasis. 2018;16(5):973–983. doi:https://doi.org/10.1111/jth.13986.

17. Tomaiuolo M, Stalker TJ, Welsh JD, Diamond SL, Sinno T, Brass LF. A systems approach to hemostasis: 2. Computational analysis of molecular transport in the thrombus microenvironment. Blood. 2014;124(11):1816–1823. doi:10.1182/blood-2014-01-550343.

18. Xu S, Xu Z, Kim OV, Litvinov RI, Weisel JW, Alber M. Model predictions of deformation, embolization and permeability of partially obstructive blood clots under variable shear flow. Journal of The Royal Society Interface. 2017;14(136):20170441. doi:10.1098/rsif.2017.0441.

19. Arrarte Terreros N, Tolhuisen ML, Bennink E, de Jong HWAM, Beenen LFM, Majoie CBLM, et al. From perviousness to permeability, modelling and measuring intra-thrombus flow in acute ischemic stroke. Journal of Biomechanics. 2020;111:110001. doi:https://doi.org/10.1016/j.jbiomech.2020.110001.

20. Du J, Kim D, Alhawael G, Ku DN, Fogelson AL. Clot permeability, agonist transport, and platelet binding kinetics in arterial thrombosis. Biophysical Journal. 2020;119(10):2102–2115. doi:https://doi.org/10.1016/j.bpj.2020.08.041.

21. Wufsus AR, Macera NE, Neeves KB. The hydraulic permeability of blood clots as a function of fibrin and platelet density. Biophysical Journal. 2013;104(8):1812–1823. doi:https://doi.org/10.1016/j.bpj.2013.02.055.

22. Muthard RW, Diamond SL. Blood lots are rapidly assembled hemodynamic sensors. Arteriosclerosis, Thrombosis, and Vascular Biology. 2012;32(12):2938–2945. doi:10.1161/ATVBAHA.112.300312.

23. Kim OV, Xu Z, Rosen ED, Alber MS. Fibrin networks regulate protein transport during thrombus development. PLOS Computational Biology. 2013;9(6):1–10. doi:10.1371/journal.pcbi.1003095.

24. Sinegre T, Milenkovic D, Teissandier D, Fully P, Bourdin J, Morand C, et al. Impact of epicatechin on fibrin clot structure. European Journal of Pharmacology. 2021;893:173830. doi:https://doi.org/10.1016/j.ejphar.2020.173830.

25. Kozeny J. Uber kapillare Leitung der Wasser im Boden. Royal Academy of Science, Vienna, Proc Class I. 1927;136:271–306.

26. Carman PC. Permeability of saturated sands, soils and clays. The Journal of Agricultural Science. 1939;29(2):262–273. doi:10.1017/S0021859600051789.

27. Slaboch CL, Alber MS, Rosen ED, Ovaert TC. Mechano-rheological properties of the murine thrombus determined via nanoindentation and finite element modeling. Journal of the Mechanical Behavior of Biomedical Materials. 2012;10:75–86. doi:https://doi.org/10.1016/j.jmbbm.2012.02.012.

28. Kadri OE, Chandran VD, Surblyte M, Voronov RS. In vivo measurement of blood clot mechanics from computational fluid dynamics based on intravital microscopy images. Computers in Biology and Medicine. 2019;106:1–11. doi:https://doi.org/10.1016/j.compbiomed.2019.01.001.

29. Wang W, Lindsey JP, Chen J, Diacovo TG, King MR. Analysis of early thrombus dynamics in a humanized mouse laser injury model. Biorheology. 2014;51:3–14. doi:10.3233/BIR-130648.

30. Taylor JO, Witmer KP, Neuberger T, Craven BA, Meyer RS, Deutsch S, et al. In vitro quantification of time dependent thrombus size using magnetic resonance imaging and computational simulations of thrombus surface shear stresses. Journal of Biomechanical Engineering. 2014;136(7). doi:10.1115/1.4027613.

31. Pinar IP, Arthur JF, Andrews RK, Gardiner EE, Ryan K, Carberry J. Methods to determine the Lagrangian shear experienced by platelets during thrombus growth. PLOS ONE. 2015;10(12):1–14. doi:10.1371/journal.pone.0144860.

32. Voronov RS, Stalker TJ, Brass LF, Diamond SL. Simulation of intrathrombus fluid and solute transport ssing in vivo clot structures with single platelet resolution. Annals of Biomedical Engineering. 2013;41(6):1297–1307. doi:10.1007/s10439-013-0764-z.

33. Fedorov A, Beichel R, Kalpathy-Cramer J, Finet J, Fillion-Robin JC, Pujol S, et al. 3D Slicer as an image computing platform for the Quantitative Imaging Network. Magnetic Resonance Imaging. 2012;30(9):1323–1341. doi:https://doi.org/10.1016/j.mri.2012.05.001.

34. Cignoni P, Callieri M, Corsini M, Dellepiane M, Ganovelli F, Ranzuglia G. MeshLab: an open-source mesh processing tool. In: Scarano V, Chiara RD, Erra U, editors. Eurographics Italian Chapter Conference. The Eurographics Association; 2008.

35. Geuzaine C, Remacle JF. Gmsh: A 3-D finite element mesh generator with built-in pre- and post-processing facilities. International Journal for Numerical Methods in Engineering. 2009;79(11):1309–1331. doi:https://doi.org/10.1002/nme.2579.

36. Kaviany M. Principles of heat transfer in porous media Springer. New York etc. 1991;.

37. Bear J, Bachmat Y. Introduction to modeling of transport phenomena in porous media. vol. 4. Springer Science & Business Media; 2012.

38. Demirin H, Ozhan H, Ucgun T, Celer A, Bulur S, Cil H, et al. Normal range of mean platelet volume in healthy subjects: Insight from a large epidemiologic study. Thrombosis Research. 2011;128(4):358–360. doi:https://doi.org/10.1016/j.thromres.2011.05.007.

39. Trudnowski RJ, Rico RC. Specific gravity of blood and plasma at 4 and 37 °C. Clinical Chemistry. 1974;20(5):615–616. doi:10.1093/clinchem/20.5.615.

40. Merrill EW, Pelletier GA. Viscosity of human blood: transition from Newtonian to non-Newtonian. Journal of Applied Physiology. 1967;23(2):178–182. doi:10.1152/jappl.1967.23.2.178.

41. Lee S, Jang S, Park Y. Measuring three-dimensional dynamics of platelet activation using 3-D quantitative phase imaging. bioRxiv. 2019;doi:10.1101/827436.

42. Bath P, Butterworth R. Platelet size: measurement, physiology and vascular disease. Blood coagulation & fibrinolysis : an international journal in haemostasis and thrombosis. 1996;7(2):157–161.

43. Donea J, Huerta A. 4. In: Viscous incompressible flows. John Wiley & Sons, Ltd; 2003. p. 147–208. Available from: https://onlinelibrary.wiley.com/doi/abs/10.1002/0470013826.ch4.

44. Hecht F. New development in FreeFem++. J Numer Math. 2012;20(3-4):251–265.

45. van der Vorst HA. Iterative Krylov methods for large linear systems. Cambridge Monographs on Applied and Computational Mathematics. Cambridge University Press; 2003.

46. Balay S, Abhyankar S, Adams MF, Benson S, Brown J, Brune P, et al. PETSc Web page; 2022. https://petsc.org/. xAvailable from: https://petsc.org/.

47. Sakariassen KS, Orning L, Turitto VT. The impact of blood shear rate on arterial thrombus formation. Future Science OA. 2015;1(4). doi:10.4155/fso.15.28.

48. Miyazaki Y, Nomura S, Miyake T, Kagawa H, Kitada C, Taniguchi H, et al. High shear stress can initiate both platelet aggregation and shedding of procoagulant containing microparticles. Blood. 1996;88(9):3456–3464. doi:https://doi.org/10.1182/blood.V88.9.3456.bloodjournal8893456.

49. Reininger AJ, Heijnen HFG, Schumann H, Specht HM, Schramm W, Ruggeri ZM. Mechanism of platelet adhesion to von Willebrand factor and microparticle formation under high shear stress. Blood. 2006;107(9):3537–3545. doi:https://doi.org/10.1182/blood-2005-02-0618.

50. Varga-Szabo D, Braun A, Nieswandt B. Calcium signaling in platelets. Journal of thrombosis and haemostasis : JTH. 2009;7(7):1057–1066. doi:10.1111/j.1538-7836.2009.03455.x.

51. Rickles FR, Falanga A. Molecular basis for the relationship between thrombosis and cancer. Thrombosis Research. 2001;102(6):V215–V224. doi:https://doi.org/10.1016/S0049-3848(01)00285-7.

52. Folie BJ, McIntire LV. Mathematical analysis of mural thrombogenesis. Concentration profiles of platelet-activating agents and effects of viscous shear flow. Biophysical Journal. 1989;56(6):1121–1141. doi:https://doi.org/10.1016/S0006-3495(89)82760-2.

53. Levine HA, McGee MP, Serna S. Diffusion and reaction in the cell glycocalyx and the extracellular matrix. Journal of Mathematical Biology. 2010;60(1):1–26. doi:10.1007/s00285-009-0254-y.

54. Teeraratkul C, Mukherjee D. Microstructure aware modeling of biochemical transport in arterial blood clots. Journal of Biomechanics. 2021;127:110692. doi:https://doi.org/10.1016/j.jbiomech.2021.110692.

55. Sadler JE. Biochemistry and genetics of von Willebrand factor. Annual Review of Biochemistry. 1998;67(1):395–424. doi:10.1146/annurev.biochem.67.1.395.

56. Savage B, Almus-Jacobs F, Ruggeri ZM. Specific synergy of multiple substrate–receptor interactions in platelet thrombus formation under flow. Cell. 1998;94(5):657–666. doi:https://doi.org/10.1016/S0092-8674(00)81607-4.

57. Ruggeri ZM, Orje JN, Habermann R, Federici AB, Reininger AJ. Activation-independent platelet adhesion and aggregation under elevated shear stress. Blood. 2006;108(6):1903–1910. doi:10.1182/blood-2006-04-011551.

58. Zhang X, Halvorsen K, Zhang CZ, Wong WP, Springer TA. Mechanoenzymatic cleavage of the ultralarge vascular protein von Willebrand factor. Science. 2009;324(5932):1330–1334. doi:10.1126/science.1170905.

59. Sing CE, Alexander-Katz A. Elongational flow induces the unfolding of von Willebrand factor at physiological flow rates. Biophysical Journal. 2010;98(9):L35–L37. doi:https://doi.org/10.1016/j.bpj.2010.01.032.

60. Schneider SW, Nuschele S, Wixforth A, Gorzelanny C, Alexander-Katz A, Netz RR, et al. Shear-induced unfolding triggers adhesion of von Willebrand factor fibers. Proceedings of the National Academy of Sciences. 2007;104(19):7899–7903. doi:10.1073/pnas.0608422104.

61. van Rooij BJM, Závodszky G, Hoekstra AG, Ku DN. Biorheology of occlusive thrombi formation under high shear: in vitro growth and shrinkage. Scientific Reports. 2020;10(1):18604. doi:10.1038/s41598-020-74518-7.

62. Petrie CJS. Extensional viscosity: A critical discussion. Journal of Non-Newtonian Fluid Mechanics. 2006;137(1):15–23. doi:https://doi.org/10.1016/j.jnnfm.2006.01.011.

63. Spieker CJ, Závodszky G, Mouriaux C, van der Kolk M, Gachet C, Mangin PH, et al. The effects of micro-vessel curvature induced elongational flows on platelet adhesion. Annals of Biomedical Engineering. 2021;49(12):3609–3620. doi:10.1007/s10439-021-02870-4.

64. Wang Y, Morabito M, Zhang XF, Webb E, Oztekin A, Cheng X. Shear-induced extensional response behaviors of tethered von Willebrand factor. Biophysical Journal. 2019;116(11):2092–2102. doi:https://doi.org/10.1016/j.bpj.2019.04.025.

65. Kania S, Oztekin A, Cheng X, Zhang XF, Webb E. Predicting pathological von Willebrand factor unraveling in elongational flow. Biophysical Journal. 2021;120(10):1903–1915. doi:https://doi.org/10.1016/j.bpj.2021.03.008.

66. Kim S, Ong PK, Yalcin O, Intaglietta M, Johnson PC. The cell-free layer in microvascular blood flow. Biorheology. 2009;46:181–189. doi:10.3233/BIR-2009-0530.

67. Fedosov DA, Caswell B, Popel AS, Karniadakis GE. Blood flow and cell-free layer in microvessels. Microcirculation. 2010;17(8):615–628. doi:https://doi.org/10.1111/j.1549-8719.2010.00056.x.

68. Zhang J, Johnson PC, Popel AS. Effects of erythrocyte deformability and aggregation on the cell free layer and apparent viscosity of microscopic blood flows. Microvascular Research. 2009;77(3):265–272. doi:https://doi.org/10.1016/j.mvr.2009.01.010.

69. Nader E, Skinner S, Romana M, Fort R, Lemonne N, Guillot N, et al. Blood rheology: key parameters, impact on blood flow, role in sickle cell disease and effects of exercise. Frontiers in Physiology. 2019;10. doi:10.3389/fphys.2019.01329.

70. Windberger U, Bartholovitsch A, Plasenzotti R, Korak KJ, Heinze G. Whole lood viscosity, plasma viscosity and erythrocyte aggregation in nine mammalian species: reference values and comparison of data. Experimental Physiology. 2003;88(3):431–440. doi:https://doi.org/10.1113/eph8802496.

