## Supplementary material for "Image-based Flow Simulation of Platelet Aggregates under Different Shear Rates": S1 File

### Platelet density measurement

In order to estimate the platelet density inside the aggregates, we used a reconstruct-volume method by manually segmenting single platelets from the aggregate in 3D Slicer. An area in which the platelets can be clearly identified in the aggregate was defined. The platelets and the aggregate in the area and its upper and lower slices were segmented manually as shown in Fig 1. A volume was therefore computed by considering several consecutive slicers. Then by analyzing the volumes of the segmented platelets and segmented aggregate, the ratio, which represents the volume fraction of platelets was computed. Simultaneously, the average fluorescence intensity of this area and its upper and lower slices were calculated. Three such areas were evaluated under each WSR, and by linear regression, a relation of fluorescence intensity and density of the entire aggregate was obtained. However, for the images captured under a higher WSR, the platelets cluster densely therefore it is hard to find out any area where platelets could be manually segmented.

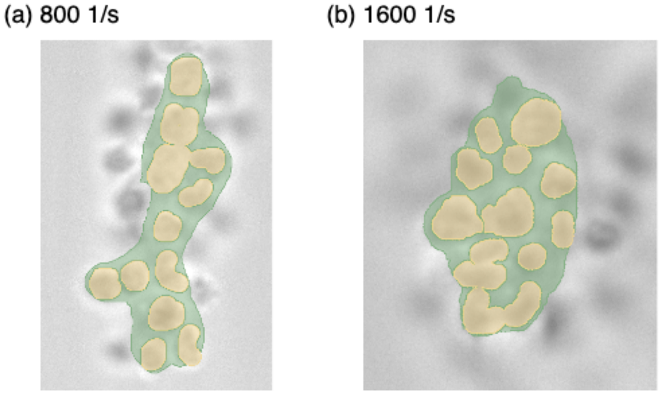


**Fig 1. 3D Slicer measurement examples.** Two examples of 3D Slicer segmentation for (a) 800 $s^{-1}$ and (b) 1600 $s^{-1}$ WSRs. The green area is the aggregate, and the yellow area represents the single platelets.

To resolve this issue, the density was estimated by computing an area ratio that could represent an 1 $\mu m$ depth volume in the 2D images. This measurement consists of the following steps. First, a fixed-radius circle was defined and assigned to five different places in non-labelled images representing possible different densities of aggregates (see Fig 2). The platelets (cross-sections) in the circle were manually counted and we assumed the cross-section of platelets was a circle with an average area of $3.1416 \mu m^2$. The density was subsequently computed by the ratio between the occupied area of platelets (the number of the platelet counted in the circle multiplied by the average area of the platelet cross-section) and the area of the defined circle. Since the distance between two slices is 1 $\mu m$, and the radius of a platelet is also 1 $\mu m$, this area ratio was applied as the density of platelets in a volume with 1 $\mu m$ depth. We also validated the estimations of platelet density of the aggregate formed under 1600 $s^{-1}$ using the reconstruct-volume method on the same circles and the average error was 0.63%. Linear regression was subsequently applied to estimate the mapping between fluorescence intensity and density of the entire aggregates based on the data of the defined circle. In this work, this method was applied and the final results are demonstrated in Fig 3 in the main article.

Based on the average platelet density inside the aggregates calculated from the method above, the number of platelets inside the aggregates could be estimated by the ratio of the non-void volume inside the aggregate and the single platelet volume. Here we assumed the volume of single platelet was 3.25 $\mu m^3$ [1]. The

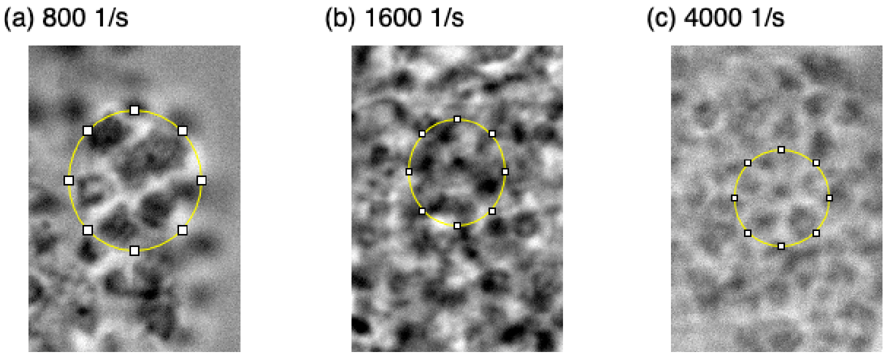


**Fig 2. Manually counting measurement examples.** Three examples of density measurement under (a) 800 $s^{-1}$, (b) 1600 $s^{-1}$ and (c) 4000 $s^{-1}$ WSRs. The platelets inside the yellow circle were counted. Here, 7 platelets were counted in each of (a), (b) and (c).

estimated numbers of platelet are around 1800, 4300 and 6200 for the platelet aggregates formed under 800 $s^{-1}$, 1600 $s^{-1}$ and 4000 $s^{-1}$ WSRs, respectively.

### References

[1] Zavodszky, G., Van Rooij, B., Azizi, V., Alowayyed, S. & Hoekstra, A. Hemocell: a high-performance microscopic cellular library. *Procedia Computer Science*. **108** pp. 159-165 (2017), https://www.sciencedirect.com/science/article/pii/S1877050917306245, International Conference on Computational Science, ICCS 2017, 12-14 June 2017, Zurich, Switzerland
