## Supplementary figures and images for "Image-based Flow Simulation of Platelet Aggregates under Different Shear Rates"

### S1 Fig

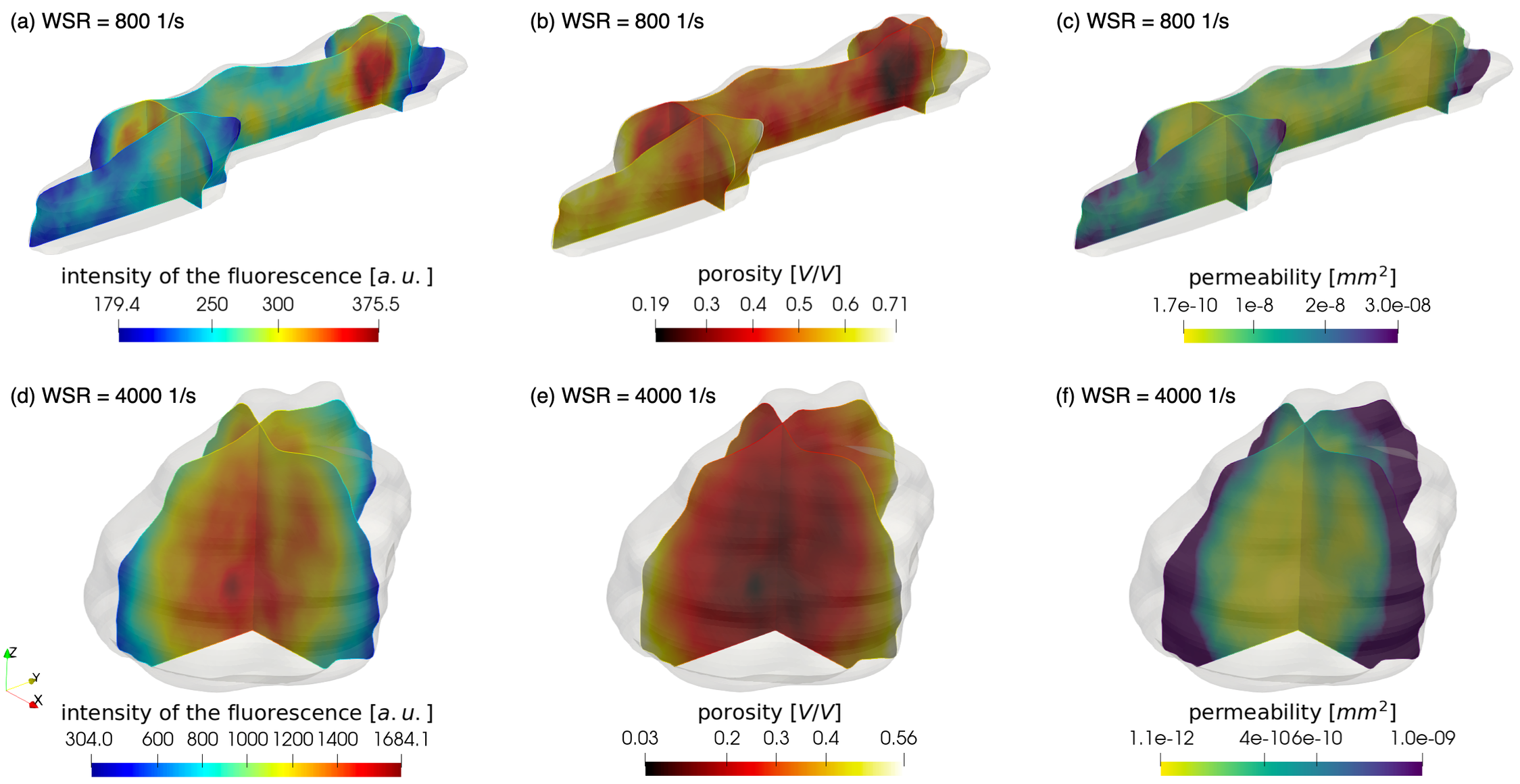

### S2 Fig

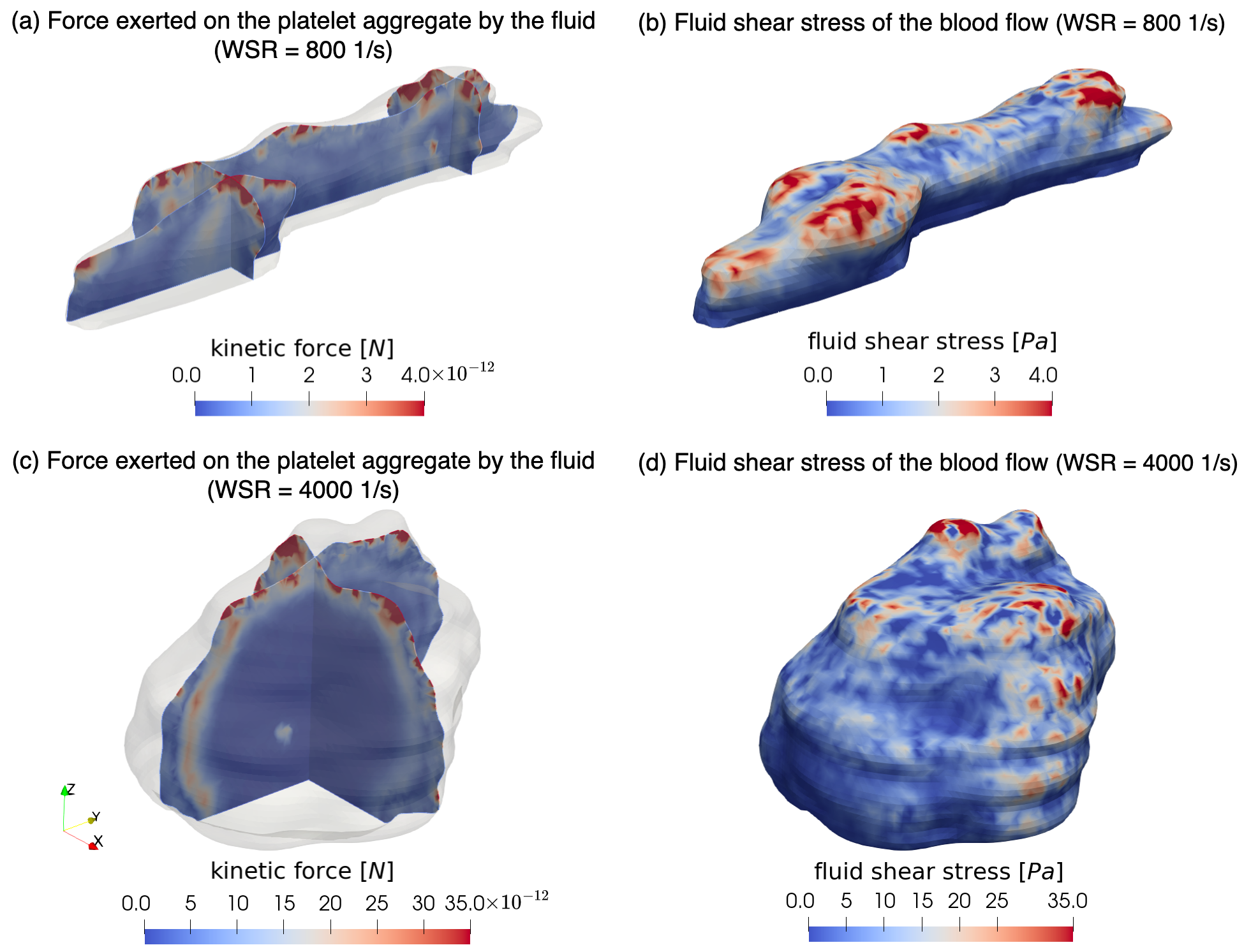

### S3 Fig

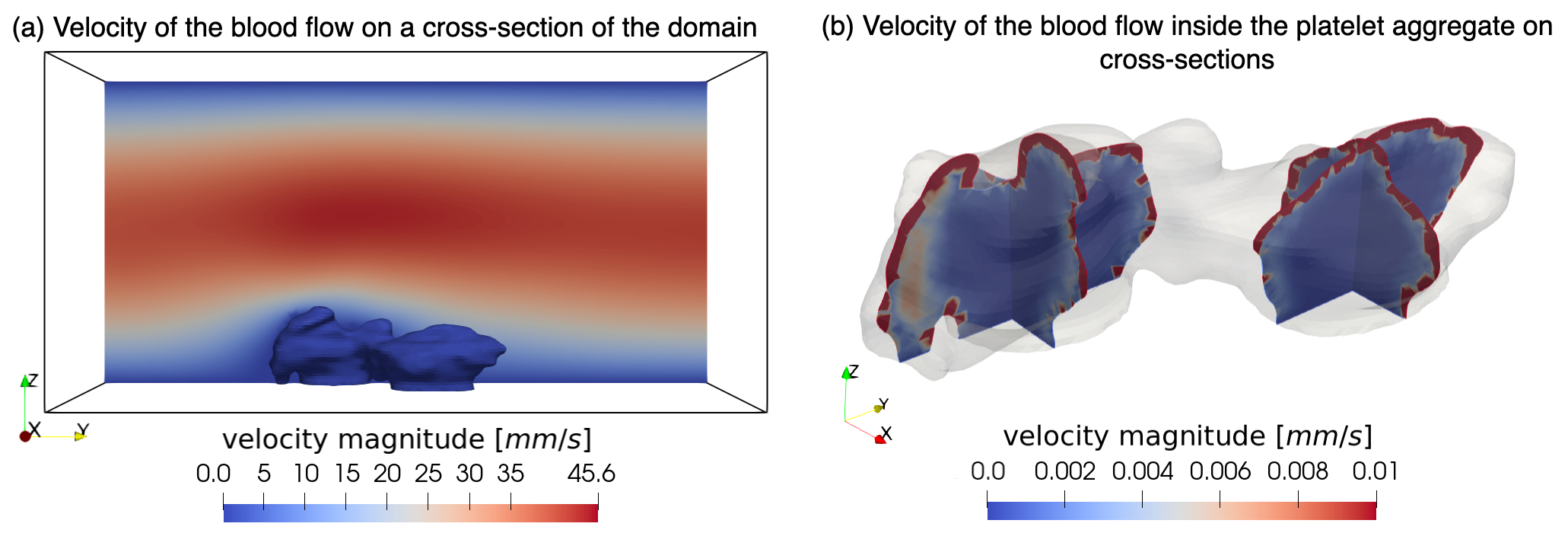
